# Computational model demonstrates that Ndc80-associated proteins strengthen kinetochore-microtubule attachments in metaphase

**DOI:** 10.1101/678789

**Authors:** Samuel Campbell, Mohammed Abdullahel Amin, Dileep Varma, Tamara Carla Bidone

**Affiliations:** Department of Bioengineering, University of Utah, Salt Lake City, UT, 84112; Department of Cell and Molecular Biology, Feinberg School of Medicine, Northwestern University, Chicago, IL; Scientific Computing and Imaging Institute, University of Utah, Salt Lake City, UT 84112; School of Computing, University of Utah, Salt Lake City, UT 84112

## Abstract

Chromosome segregation is mediated by spindle microtubules that attach to kinetochore via dynamic protein complexes, such as Ndc80, Ska, Cdt1 and ch-TOG during mitotic metaphase. While experimental studies have previously shown that these proteins and protein complexes are all essential for maintaining a stable kinetochore-microtubule interface, their exact roles in this process remains elusive. In this study, we employed experimental and computational methods in order to characterize how these proteins can strengthen kMT attachments in both non load-bearing and load-bearing conditions, typical of prometaphase and metaphase, respectively. Immunofluorescence staining of Hela cells showed that the levels of Ska and Cdt1 significantly increase from prometaphase to metaphase. Our new computational model showed that, by incorporating binding and unbinding of each protein complex, coupled with a biased diffusion mechanism, the displacement of a possible complex formed by Ndc80-Ska-Cdt1 is significantly higher than that formed by Ndc80 alone or Ndc80-Ska. In addition, when we use Ndc80/ch-TOG in the model, rupture force and time of attachment of the kMT interface increases. These results support the hypothesis that Ndc80-associated proteins strengthen kMT attachments and that it is the interplay between kMT protein complexes in metaphase that ensures stable attachments.

## Introduction

During cell division, stable attachments between kinetochores of the chromosomes and spindle microtubules are essential for ensuring accurate segregation of the genetic material. In metaphase and anaphase, microtubules undergo structural changes via lengthening and shortening of tubulin protofilaments that generate forces at the kinetochore-microtubule (kMT) interface (Jennifer G. DeLuca & Musacchio, 2012; Jeyaprakash et al., 2012). In metaphase, the G-protein pathway activates pulling forces to align chromosomes at the metaphase plate (Grill, Gönczy, Stelzer, & Hyman, 2001; Grill, Howard, Schäffer, Stelzer, & Hyman, 2003). Directional motility of chromosomes driven by kinetochore motors such as dynein and CENP-E and chromokinesins that bind to chromosome arms also contribute critically to chromosome congression and alignment (Maiato, Gomes, Sousa, & Barisic, 2017). Remarkably, kMT attachments remain robust and withstand tension to avoid erroneous segregation. However, how kinetochores remain associated to dynamic microtubule ends under tension during anaphase and metaphase remains not fully understood.

In metaphase, kMT attachments require several proteins and protein complexes, including Ndc80, Ska1, Cdt1 and ch-TOG (G. Alushin & Nogales, 2011; Jennifer G. DeLuca & Musacchio, 2012; Lampert & Westermann, 2011; Takeuchi & Fukagawa, 2012). These proteins factors can broadly be generalized as microtubule-associated proteins (MAPs) at kinetochores that possess the ability to bind to spindle microtubules during metaphase. They are able to undergo dynamic cycles of binding, diffusion and unbinding along tubulin protofilaments (Agarwal et al., 2018; G. M. Alushin et al., 2010; Cooper & Wordeman, 2009; Gestaut et al., 2008; Jeyaprakash et al., 2012; Powers et al., 2009; Spittle, Charrasse, Larroque, & Cassimeris, 2000; Westermann et al., 2006). The Ndc80 complex is the major protein component of the kMT interface and creates a direct link between kinetochores and microtubules (Cheeseman, Chappie, Wilson-Kubalek, & Desai, 2006). Affinity and avidity regulate Ndc80’s binding rates, while Aurora B kinase phosphorylation regulates its unbinding rates, resulting in a biased diffusion mechanism with diffusion constants in the range of *D* = 0.018 −0.03 μm^2^·s^−1^ (Powers et al., 2009; Umbreit et al., 2012). Ska1 and Cdt1 bind to Ndc80 and form load-bearing bridges between Ndc80 and tubulin protofilaments (Cheeseman et al., 2001; Davis et al., 2018; Hanisch, Silljé, & Nigg, 2006; Schmidt et al., 2012; Welburn et al., 2009). Ska1 and Cdt1 diffuse along microtubules with average diffusion rates *D* = 0. 09 μm^2^·s^−1^ and *D* = 0.16 μm^2^·s^−1^, respectively (Agarwal et al., 2018; Schmidt et al., 2012). The Ska complex strengthen Ndc80 ability to track microtubules ends, by also dampening chromosome motion to promote accurate segregation (Cheerambathur et al., 2017; Schmidt et al., 2012). Cdt1 binding affinity to microtubules is also controlled by Aurora B kinase phosphorylation (Agarwal et al., 2018; Dileep Varma et al., 2012). An additional protein, ch-TOG, a non-motor protein complex that organizes spindle poles, has been recently found to be critical for load-bearing kMT attachments (Gergely, Draviam, & Raff, 2003). Experiments using optical traps showed that ch-TOG’s activity is tension-dependent and stabilizes kMT attachments (Miller, Asbury, & Biggins, 2016).

Previous studies have collectively led to the idea that Ndc80, Ska, Cdt1 and ch-TOG are all critical for stabilizing kMT attachments during metaphase. However, we still lack a clear understanding of the precise roles played by these individual factors and/or how their interplay contributes to kMT attachment stability. An interesting hypothesis is that these proteins directly strengthen the kinetochore-microtubule interface by forming additional connections between the Ndc80 complex and the microtubule (Davis et al., 2018). However, since these proteins dynamically form and break their connections with tubulin protofilaments, while diffusing on microtubules, a synergy between their activities is likely to exist (Miller et al., 2016; Schmidt et al., 2012). Two models have been previously proposed in order to understand how multiple proteins at kMT attachments withstand tension: according to the ring model, tension from dynamic microtubules slides a ring of proteins (Koshland, Mitchison, & Kirschner, 1988); in the fibrils model, kinetochores movement is driven by tugging on tightly bound kinetochore protein fibrils (McIntosh et al., 2008). While providing valuable insights into the molecular mechanisms of multiprotein kMT attachments, these models did not directly incorporate parameters of Ndc80, Ska, Cdt1 and ch-TOG. Therefore, a detailed understanding of the individual contributions from these proteins and proteins complexes and/or interplay is currently missing.

In this work, we used high-resolution fluorescence microscopy and computational simulation approaches in order to understand the interplay between kMT proteins in prometaphase and metaphase. We found that load-bearing kMT attachments in metaphase have considerable higher level of Ska and Cdt1 with respect to non load-bearing attachments in prometaphase. In order to understand the synergy between the different kinetochore MAPs with Ndc80, we developed a new computational model. We characterized the effect of binding and unbinding from Ndc80, Ska, Cdt1 and ch-TOG on kMT attachment stability. In particular, we used the model to elucidate how binding and unbinding rates of individual protein components, in combination with a biased diffusion mechanism and tension-dependent bond lifetimes, can strengthen kMT attachments. In load-bearing conditions, by increasing the force across the kinetochore-microtubule interface, we also emulated detachment of kMT interfaces. The model showed that combining Ndc80, Ska and Cdt1 enhances kMT attachment strength with respect to individual proteins. Using Ndc80/ch-TOG also strengthens the complex with respect to both Ndc80 in isolation or Ndc80-Ska-Cdt1. Taken together, our results provide important mechanistic insights into how critical kMT proteins interplay to withstand tension and ensure accurate chromosome segregation.

## Results

### Higher levels of Ska and Cdtl at metaphase as compared to prometaphase kinetochores

We first aimed to address the roles played by kinetochore microtubule-binding proteins Ska and Cdt1 towards the formation of load-bearing attachments by the Ndc80 complex. To analyze the difference in kinetochore localization between a non-load bearing (kinetochore pairs not attached to spindle microtubules) and a load bearing (bioriented kinetochores attached to spindle microtubules from opposite spindle poles) state, we assayed the kinetochore levels of Ska and Cdt1 in prometaphase and metaphase mitotic cells respectively. We found that there is a considerable increase in kinetochore levels of both these proteins from prometaphase to metaphase (Fig. 1 C-F; (Agarwal et al., 2018; Chan, Jeyaprakash, Nigg, & Santamaria, 2012; Hanisch et al., 2006; Dileep Varma et al., 2012)). There was a robust 4-fold increase in Ska levels in metaphase compared to prometaphase kinetochores (Fig. 1 C, D; (Chan et al., 2012; Hanisch et al., 2006). The increase in levels of Cdt1 at metaphase was consistent but marginal as compared to prometaphase kinetochores (Fig. 1 E, F). The kinetochore levels of the Hec1 subunit of the Ndc80 complex, as expected, did not vary much between prometaphase and metaphase (Fig. 1 A, B; (Gascoigne & Cheeseman, 2013; Lin, Chen, Wu, & Lee, 2006)).

**Figure 1:**
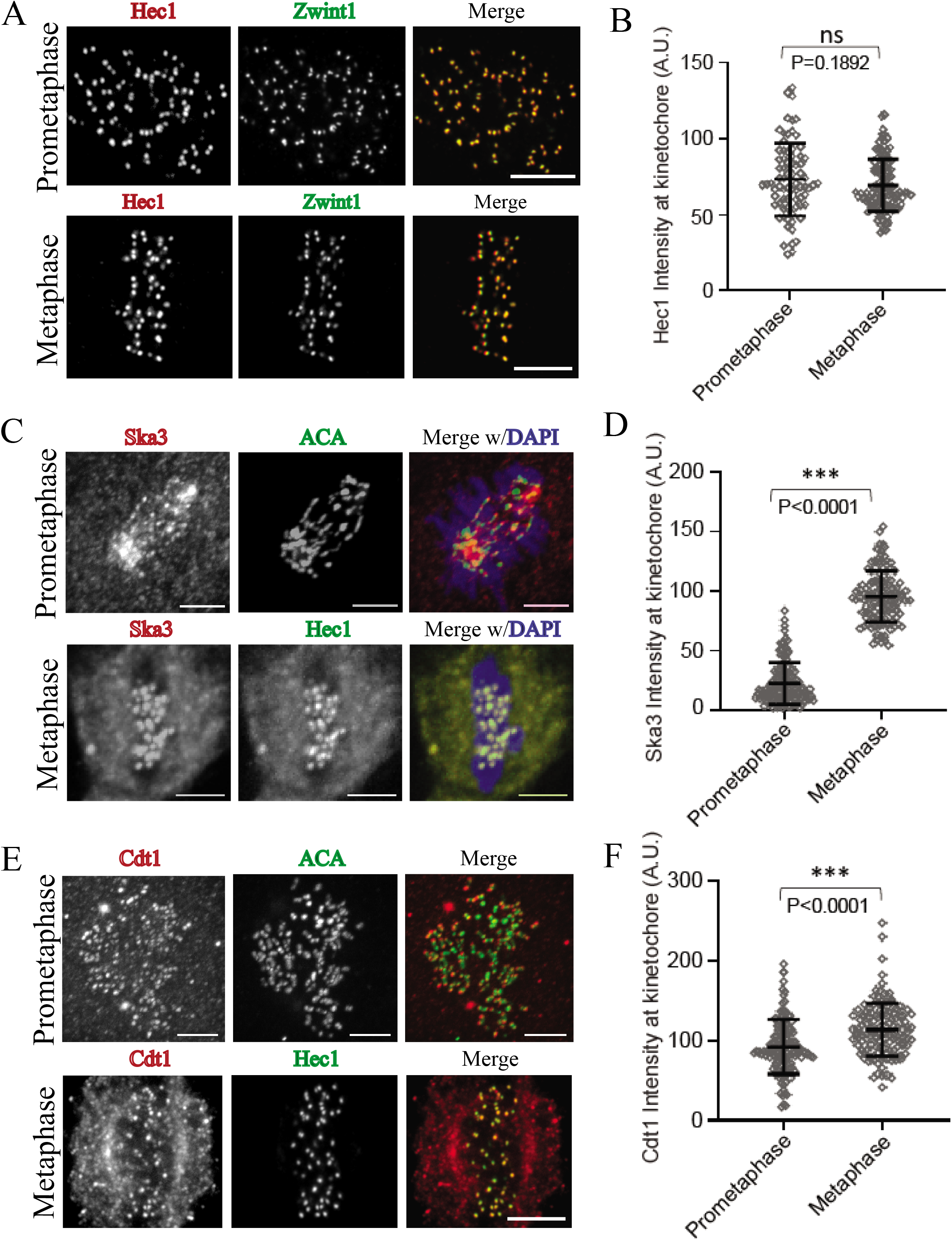
Kinetochore localization of microtubule-binding proteins that are required for stabilizing end-on kinetochore-microtubule attachments in fixed mitotic HeLa cells. (A) Immunofluorescence staining of mitotic cells for the Hec1 subunit of the Ndc80 complex shown alongside another kinetochore marker, Zwint1 in either prometaphase (top panel) or metaphase (bottom panel). (B) Comparative quantification of Hec1 intensity at kinetochores in prometaphase vs metaphase cells from A. n = 75 from 5 cells. (C) Immunofluorescence staining of mitotic cells for the Ska3 subunit of the Ska complex shown alongside kinetochore markers, anti-CREST antiserum (ACA) in prometaphase (top panel) or Hec1 in metaphase (bottom panel), as indicated. (D) Comparative quantification of Ska3 intensity in prometaphase *vs* metaphase cells from C. n=120 from 5 cells. (E) Immunofluorescence staining of mitotic cells for Cdt1 shown alongside kinetochore markers, anti-CREST antiserum (ACA) in prometaphase (top panel) or Hec1 in metaphase (bottom panel), as indicated. (F) Comparative quantification of Cdt1 intensity in prometaphase *vs* metaphase cells from E. n=120 from 5 cells.

### Model calibration

We calibrated the kinetic Monte Carlo model (Fig. 2A) based upon experimentally detected parameters for binding, unbinding and diffusion of Ndc80, Ska and Cdt1 (Agarwal et al., 2018; Schmidt et al., 2012; Umbreit et al., 2012). Initially, each protein was simulated in isolation. In the parameter space defined by *k*_on_ and *k*_off_, we identified diffusion constants, *D*, for Ndc80, Ska and Cdt1 (Fig. 2B) and compared them with those extracted from previous *in vitro* experiments. When a binding rate was not available from experiments, we identified the *D* corresponding to the known unbinding rate and used the model to predict the binding rate. For example, for Ndc80, using *k*_off_ = 0.21 s^−1^ and diffusion coefficient in the range *D* = 0.018 −0.03 μm^2^·s^−1^, as extracted from previous Total Internal Fluorescence Microscopy (TIR-FM) experiments (Schmidt et al., 2012; Umbreit et al., 2012), we found *k*_on_ = 0.03 s^−1^. For Ska, we used unbinding rate *k*_off_ = 0.15 s^−1^, binding rate *k*_off_ = 0.16 s^−1^, and diffusion coefficient *D* = 0.09 μm^2^·s^−1^ (Schmidt et al., 2012). For Cdt1, higher binding and unbinding rates with respect to both Ndc80 and Ska were used: *k*_off_ = 0.4 s^−1^ and *k*_on_ = 0.2 s^−1^. Accordingly, Cdt1 resulted in a higher average diffusion coefficient: *D* = 0.16 μm^2^/s (Fig. 2B), consistent with experimental data (Agarwal et al., 2018).

**Figure 2.**
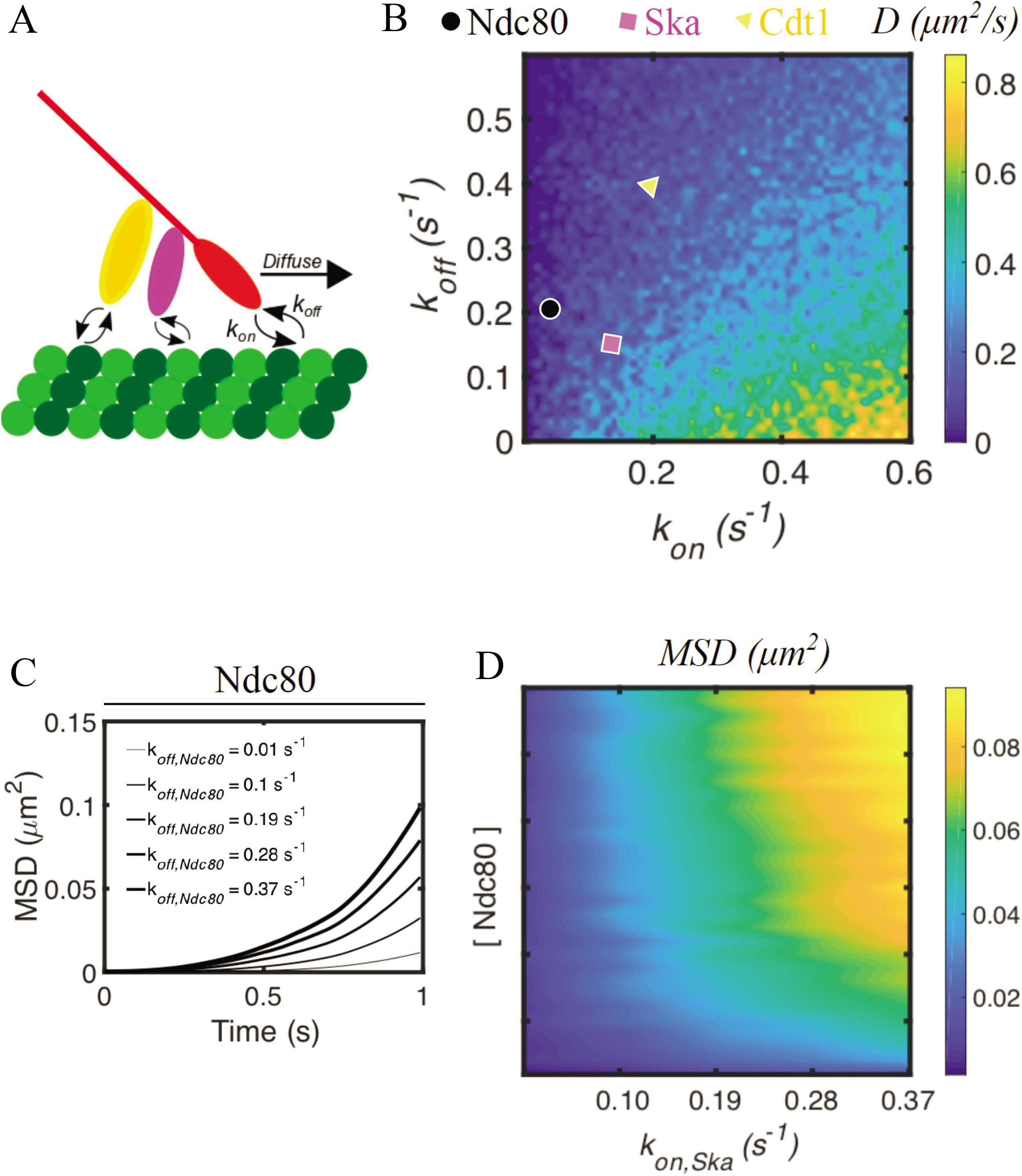
Computational model of kMT attachments incorporating the Ndc80-accessory proteins. (A) Schematics of the computational model, incorporating binding, unbinding and biased diffusion of Ndc80-associated proteins Ska and Cdt1. (B) Heatmap showing the diffusion coefficient of a protein complex undergoing biased diffusion as a function of *k_on_* and *k_off_*. Parameter values for Ndc80, Ska and Cdt1 are extracted or evaluated from previous *in vitro* characterization of the complex dynamics on immobilized microtubules. (C) Mean square displacement of Ndc80 during 1 s of simulation using different values of *k_off_*, in order to mimic the effect of phosphomimetic Ndc80 mutants. (D) Heat map representing average Mean Square displacement (MSD) of the Ndc80-Ska complex, at varying concentrations of Ndc80 and by systematically changing Ska binding rate,

### Model validation

We tested if the predictive performance of our model deteriorates substantially when applied to data that were not used in the calibration. For this, we used systems of multiple proteins and evaluated kMT attachment stability. We used the model to simulate: (*i*) Ndc80 phosphorylation (Zaytsev et al., 2015); (*ii*) Ska binding microtubules without Ndc80 (Schmidt et al., 2012). Since *k*_on_ for different numbers of Ndc80 phosphomimetic substitutions does not substantially differ (Zaytsev et al., 2015), in order to mimic Ndc80 phosphorylation, we systematically increased Ndc80 *k*_off_ and evaluated the mean square displacement, MSD, of the complex. For 1 s of simulation, increasing Ndc80 *k*_off_ from 0.01 s^−1^ to 0.37 s^−1^ increases MSD by about ten-fold, from 0. 01 μm^2^ to 0.1 μm^2^ (Fig. 2C). This result is quantitatively consistent with data from TIR-FM analysis of Ndc80 phosphomimetic mutants (Zaytsev et al., 2015).

Ska binding rate depends upon whether the protein is in isolation or in combination with Ndc80, increasing from 0.02 s^−1^ to 0.16 s^−1^, when forming a complex (Schmidt et al., 2012). Therefore, we evaluated the accuracy of our model by varying Ndc80 concentration and Ska *k*_on_. For *k*_on_ > 0.1s^−1^, MSD of the Ndc80-Ska complex increases in proportion to Ndc80 concentration, by about 8-fold (Fig. 2D), as in previous experiments (Zaytsev et al., 2015). Without Ndc80, variations of *k*_on,Ska_ minimally affect MSD. Co-sedimentation assays had previously also shown that Ndc80 increases ska affinity in a dose-dependent manner, up to 8-fold (Zaytsev et al., 2015), consistent with our modeling results.

### Binding and unbinding of Ndc80-associated proteins stabilizes kMT connections in non load-bearing conditions

In order to gain insights into the processivity of a possible complex formed by Ndc80, Ska and Cdt1, we next used the model in order to test how variations of each component’s *k_off_* reflect on the displacement of Ndc80-Ska-Cdt1. With Ndc80 only, the complex moves about 6 μm in 2 minutes; using Ndc80-Ska and Ndc80-Ska-Cdt1, the complex moves about 28 and 33 μm, respectively, in the same amount of time (Fig. 3A). The higher binding rate and smaller unbinding rate of Ska with respect to Ndc80 (Fig. 2B) significantly increase the displacement of Ndc80-Ska with respect to only Ndc80 (Fig. 3A). These results are qualitatively consistent with analysis from TIR-FM and co-sedimentation assays of Ndc80+Ska (Schmidt et al., 2012) and with *in vivo* analysis of Cdt1 in Hela cells (Agarwal et al., 2018; Dileep Varma et al., 2012), reporting that both Ska and Cdt1 stabilize kMT attachments. Results from TIR-FM had observed a 1.5-fold change in fluorescence intensity between Ndc80 alone and Ndc80-Ska (Schmidt et al., 2012). The difference in displacement between Ndc80 and Ndc80-Ska from our model is about 4-fold (Fig. 3A). A possible explanation for this difference is that the kinetic rates of Ska and Cdt1 are affected by loads and this can lead to smaller overall displacements. A similar effect on kinetic rates is induced by Aurora B kinase phosphorylation of the Ndc80 complex (Zaytsev et al., 2015).

**Figure 3.**
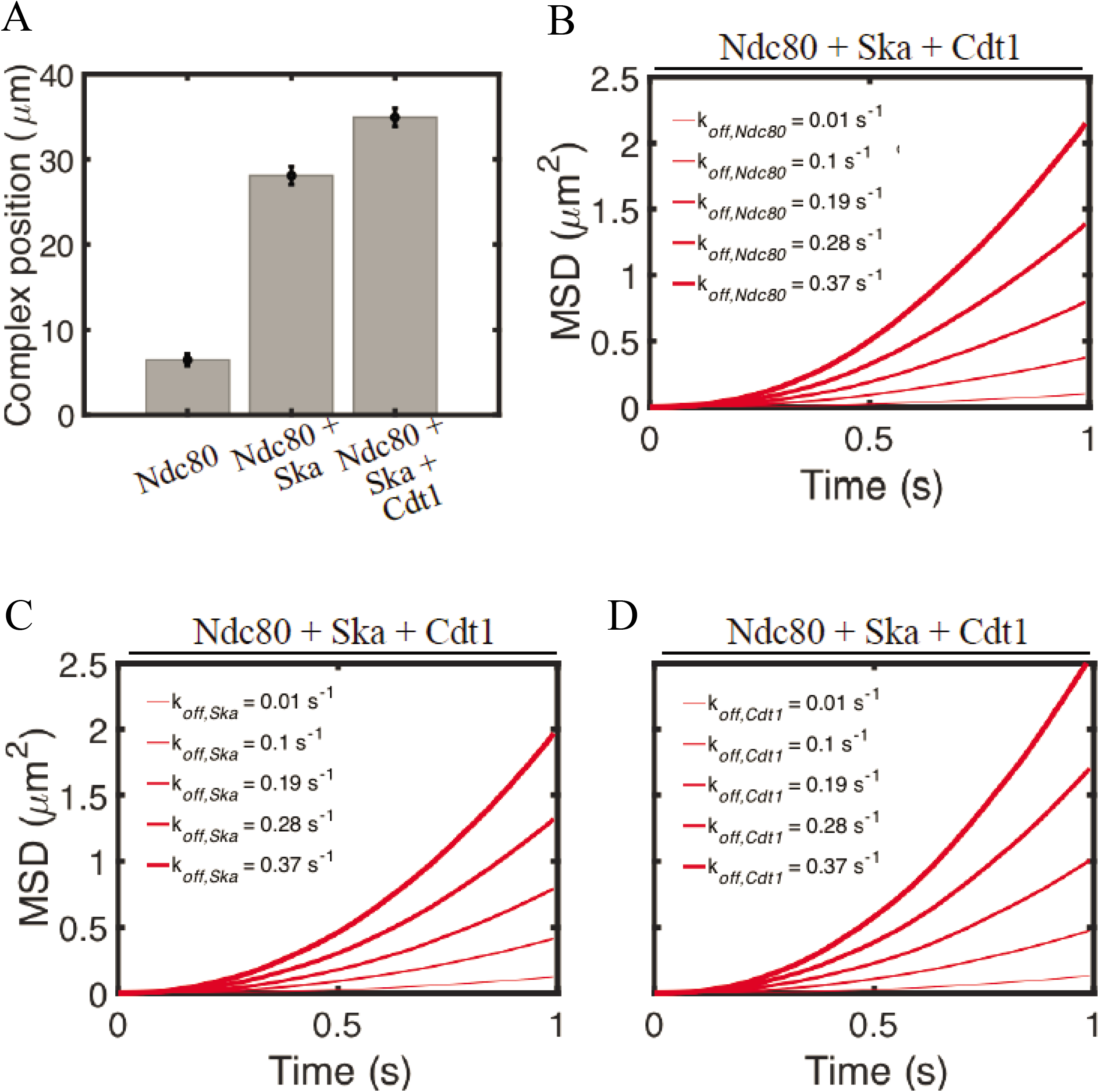
The interplay between Ndc80, Ska and Cdt1 to ensure stable kinetochore-microtubule attachment in non load-bering conditions. (A) Average position of the Ndc80 complex at 120 seconds of simulations. Error bars denote standard deviation from the mean. Data are computed from 50 independent runs. Mean square displacement (MSD) of the Ndc80 complex, including Ska and Cdt1, during 1 s of simulations, for different values of (B) *k_off,Ndc80_*, (C) *k_off,ska_* or (D) *k_off,cdt1_*.

In order to gain more insights into the effect of Aurora B on the processivity of a possible complex formed by Ndc80, Ska and Cdt1, we next tested how variations of each component’s *k_off_* reflect on the displacement of the complex Ndc80-Ska-Cdt1. By systematically increasing either Ndc80 or Ska koff, from 0.01 s^−1^ to 0.37 s^−1^, an increase in the complex MSD is observed, from about 0.2 μm^2^ to 2 μm^2^, in 1 s of simulation (Fig. 3B-C). Increasing Cdt1 *k_off_* within the same range enhances the complex MSD up to 2.5 μm^2^ in 1 s (Fig. 3D). This result indicates that phosphorylation of Cdt1 can alter the complex processivity more than Ska or Ndc80 phosphorylation.

### Role of Ndc80, Ska and Cdt1 in kMT attachment in load-bearing conditions

Next, we wanted to understand the properties of the various kinetochore MAPs in assisting the Ndc80 complex to form load-bearing kMT attachments. We first carried out immunofluorescence staining experiments to localize Ska and Cdt1 in mitotic metaphase HeLa cells. Contrary to the observed kinetochore staining in paraformaldehyde fixed cells, when we fixed the cells with methanol, we were able to discern clear mitotic spindle microtubule and spindle pole staining for both Ska 3 (top panel) and Cdt1 (bottom panel) (Fig. 4A, (Agarwal et al., 2018; Hanisch et al., 2006)). Clear Ska and Cdt1 staining on individual microtubules, however, was often difficult to discern (data not shown).

**Figure 4.**
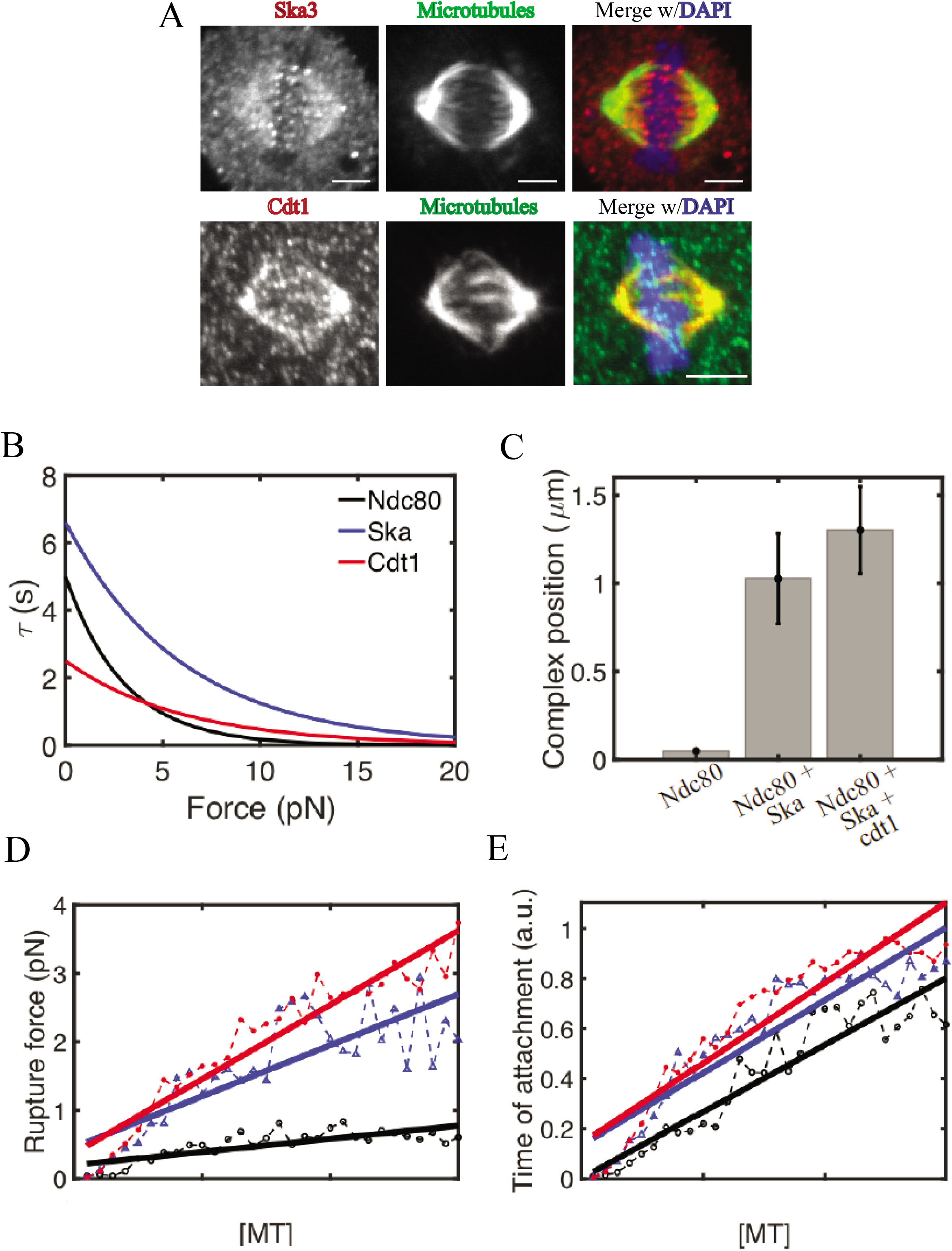
Ndc80, Ska and Cdt1 increase kinetochore-microtubule rupture force and attachment time in load-bearing conditions. (A) Spindle microtubules localization of the Ska3 subunit of the Ska complex (red, top panel) or Cdt1 (red, bottom panel) in methanol-fixed (bottom) metaphase HeLa cells. Microtubules are in green while DAPI/chromosomes are in blue. (B) Implemented bond lifetimes for Ndc80, Ska, and Cdt1. (B) Average displacement of the Ndc80 complex at 120 seconds of simulations. Error bars denote standard deviation from the mean. Data are computed from 100 independent runs. (C) Average rupture force for Ndc80, Ndc80+Ska, and Ndc80+Ska+Cdt1 versus microtubule concentration. Dashed lines indicate simulation data; solid lines represent linear fitting. (D) Average attachment lifetime for Ndc80, Ndc80+Ska, and Ndc80+Ska+Cdt1 versus microtubule concentration. Dashed lines indicate simulation data; solid lines represent linear fitting. All reported data are averages from 100 independent runs.

In order to test the effect of physiologically relevant loads on the stability of kMT attachments, we incorporated in the model explicit forces acting on bound MAPs. The model assumes that Ndc80, Ska and Cdt1 behave as slip bonds, and therefore have unbinding rates depending on the tension on the bond (Fig. 4B). Unloaded bond lifetimes for Ndc80, Ska and Cdt1 were given by the inverse of the corresponding binding rate at zero force (Fig. 2B): *τ_Ndc80_* = 4.7 s; *τ_ska_* = 6.7 *s*; *τ*_*Cdt*1_ = 2.5 *s* (Fig. 4B). Ndc80 alone moved about 0.3 μm in 120 s, while Ndc80-Ska and Ndc80-Ska-Cdt1 moved about 1 and 1.3 μm, respectively (Fig. 4C). The rupture force, corresponding to the load under which no MAP is bound, increased in proportion to the number of simulated microtubules, for Ndc80, Ndc80-Ska and Ndc80-Ska-Cdt1. In particular, rupture force was between 0.5-1pN for Ndc80, and up 3- to 4-fold higher using Ndc80-Ska and Ndc80-Ska-Cdt1, respectively (Fig. 4D). Similarly, time of attachment, corresponding to total time during which the complex is bound before rupture, increased with microtubule concentration (Fig. 4E). This time was lower using Ndc80 than Ndc80-Ska-Cdt1 (Fig. 4E). *In vitro* experiments based on laser trapping-based force clamps had shown that Ndc80 moves about 0.6 μm before detaching and that a tension in the range of 0.5-2.5 pN generates unbinding (Powers et al., 2009), fully consistent with our modeling results.

### Role of ch-TOG in kMT attachments using load-bearing conditions

Finally, we were interested in understanding the role of the kinetochore MAP ch-TOG in assisting the Ndc80 complex for the formation of load bearing kMT attachments. We first expressed a GFP-tagged ch-TOG construct in HeLa cells to characterize its localization in mitotic cells. Paraformaldehyde fixation revealed clear kinetochore localization of ch-TOG in metaphase cells (Fig. 5A). However, we were not able to discern a difference between the kinetochore staining of ch-TOG in prometaphase as compared to metaphase cells (data not shown). On the other hand, fixing metaphase cells with Methanol showed clearly definable mitotic spindle staining, with bright spindle pole staining and a milder kinetochore staining (Fig. 5A). This staining pattern was reminiscent of Ska and Cdt1 spindle and spindle pole staining after methanol fixation of metaphase cells.

**Figure 5.**
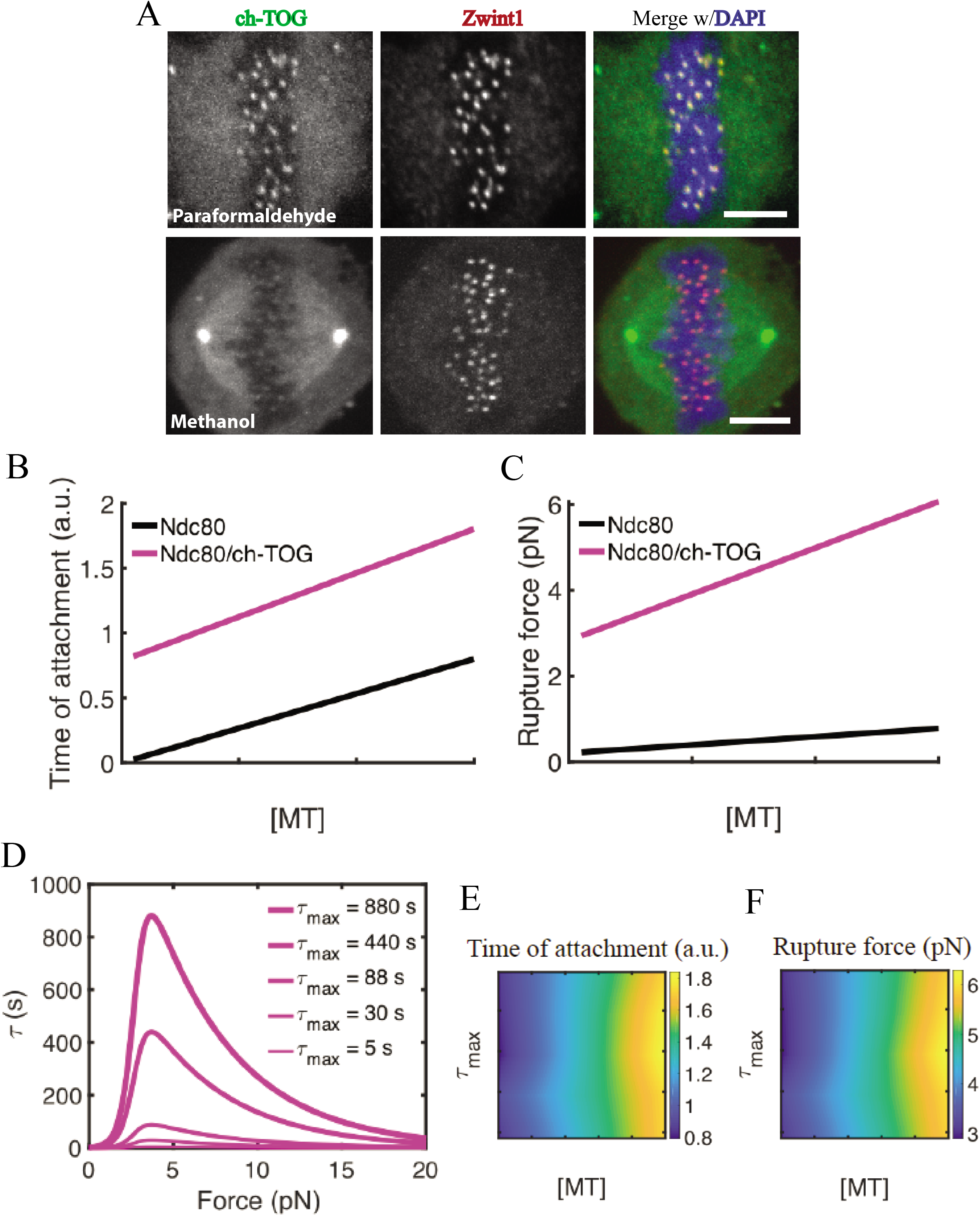
ch-TOG significantly enhances rupture force and attachment time via tension-dependent mechanisms. (A) Localization of GFP-tagged ch-TOG to kinetochores (green, top panel) in paraformaldehyde-fixed and to spindle microtubules (green, bottom panel) in methanol-fixed metaphase HeLa cells. (B) Average attachment time for Ndc80 and Ndc80+ch-TOG versus microtubule concentration. Data are extracted from linear fitting of the simulation results. Results are computed as averages from 30-60 independent runs. (C) Average rupture force for Ndc80 and Ndc80+ch-TOG versus microtubule concentration. Results are computed from 30-60 independent runs. (D) Implemented bond lifetimes for ch-TOG. Bond lifetime versus force relation for varying *τ_MAX_* are tested (E) Heatmap of average attachment time by systematically varying microtubule (MT) concentration and *τ_MAX_*. (E) Heatmap of average rupture force by systematically varying MT concentration and *τ_MAX_*.

To better understand the contribution of ch-TOG for forming robust kMT attachments, we incorporated in our model an additional protein component that directly stabilizes attachments in a force-dependent way. We used a catch bond dynamics, as detected from previous optical traps experiments (Miller et al., 2016). Over the course of the simulations, ch-TOG maintains its attachment to the kMT by sharing the total load, *F_tot_*, with the other bound proteins. With respect to Ndc80 alone, Ndc80/ch-TOG remains attached 3-fold longer (Fig. 5B). Rupture force also increases up to 5 pN, for high microtubule concentrations (Fig. 5C). Since there is no single protein experiment estimating the force versus bond lifetime of individual ch-TOG proteins, we tested how catch bond dynamics with different *τ_MAX_* (Fig. 5D) affect attachment time and rupture force at varying microtubule concentration. The model shows that by increasing *τ_MAX_* from 5 s to 880 s, time of kMT attachment and rupture force do not vary significantly (Fig. 5E-F). Instead, microtubule concentration is positively correlated with kMT attachment time and rupture force.

## Discussion

The ability of cells to separate chromosomes during mitosis is critical to several aspects of their physiology and pathology, including normal tissue development, aneuploidy and cancer. In this study, we demonstrated that a dynamic interplay between protein complexes forming links between spindle microtubules and chromosomal kinetochores is required to ensure stable attachments between the two cellular structures and ensure accurate chromosome segregation. We showed that the levels of Ska and Cdt1 increase from prometaphase to metaphase (Fig. 1), supporting the hypothesis that these Ndc80-accessory proteins functionally complement kMT attachments. We tested this hypothesis by developing a computational model of kMT attachments that incorporates binding, unbinding and biased diffusion of individual protein complexes. The model showed that accessory Ndc80-binding proteins increase kMT attachment time and rupture force in load-bearing conditions (Fig. 4C-D). In addition, incorporating in the model the contribution of ch-TOG further stabilizes kMT attachments, due to its catch bond dynamics (Fig. 5B-C). These effects result from binding, biased diffusion and tension-dependent unbinding of the protein complexes.

Previous studies of kMT attachments have shown that an increase in Ndc80 surface density results in the strengthening of kMT attachments, as more Ndc80 complexes can reach tubulin protofilaments, bind them and withstand mitotic forces (Davis et al., 2018). Accordingly, Ndc80 knockdowns *in vivo* cause severe defects in kMT attachments (J. G. DeLuca, 2004; McCleland et al., 2004; D. Varma & Salmon, 2013; Wei, Al-Bassam, & Harrison, 2007; Wigge & Kilmartin, 2001). Our model is consistent with these results showing that an increase in Ndc80 density increases the displacement of the Ndc80-Ska complex (Fig. 2D). Moreover, previous experiments have shown that Ska and Cdt1 are recruited to kinetochores by Ndc80 to provide more stability to attachments (Gaitanos et al., 2009; Jeyaprakash et al., 2012; Raaijmakers, Tanenbaum, Maia, & Medema, 2009; Welburn et al., 2009), and that ch-TOG is also needed to ensure load-bearing properties to the complex. Loss of Ska *in vivo* delays mitotic progression and it has been associated with chromosome congression failure and mitotic cell death (Gaitanos et al., 2009; Welburn et al., 2009). Accordingly, deletion of Ska prevents it from strengthening Ndc80 complex-based attachments, as revealed by approaches based on force measurements employing optical tweezers (Davis et al., 2018). Our model shows that the presence of Ska allows the complex to diffuse more than with Ndc80 in isolation, both in non load-bearing and load-bearing conditions (Fig. 3A and Fig. 4C). In addition, in load-bearing conditions, Ska enhances the force required for kMT attachment rupture of about 3-fold relative to Ndc80 alone. By adding Cdt1, a smaller, but significant enhancement for Ndc80-Ska-Cdt1 attachment strength is detected with respect to Ndc80-Ska, both in non load-bearing and load-bearing conditions (Fig. 3A and Fig. 4D). When Ch-TOG is incorporated in the model, rupture force is significantly higher that with Ndc80-Ska or Ndc80-Ska-Cdt1, owing to its catch bond dynamic (Fig. 5C). This result is consistent with findings from previous *in vitro* reconstitution systems, showing a direct role for ch-TOG in stabilizing kMT attachments via its tension-dependent bond dynamics (Miller et al., 2016).

This study contributes to the advancement of our understanding of the interplay between Ndc80-associated proteins, specifically Ska, Cdt1 and ch-TOG, in strengthening kMT attachments during metaphase. Previous theoretical models have proposed that ring, sleeves or fibrils containing ensembles of kMT proteins form multiple weak attachments with dynamic microtubules (Hill, 2006; Koshland et al., 1988; McIntosh et al., 2008). While these previous models provide insights into important mechanisms by which kMT remain attached to dynamic microtubules, they do not incorporate parameters specific to Ndc80, Ska, Cdt1 or ch-TOG, such as tension-dependent bond dynamics for these proteins. Our model directly incorporates tension-dependent bonds dynamics for Ndc80, Ska, Cdt1, and ch-TOG. By testing different conditions of proteins densities and levels of binding and unbinding under load, out model allows us to combine contributions from the various MAPs and study how the kMT complex responds to physiologically relevant tensions. In this way, our model allows us to evaluate the synergistic effect of Ndc80-associated metaphase proteins on strengthening of the kMT interface. Furthermore, different from the sleeve and ring models (Koshland et al., 1988; McIntosh et al., 2008), where the kMT proteins are rigidly connected, our model treats binding and unbinding of Ndc80-associated proteins independently. The results from our model suggest that a possible complex, Ndc80-Ska-Cdt1, in prometaphase is highly diffusive (higher levels of displacement) which agrees with the notion that kinetochore association with microtubules are dynamic and amenable to sliding motility on microtubules in non load-bearing conditions. The formation of a load-bearing network between these proteins in metaphase enables a robust, processive interaction with microtubule-ends that reduces the ability for random diffusion while at the same time enabling oscillatory behavior of kinetochores coupled to microtubule polymerization and depolymerization. This processive complex also enables the dynamic coupling of Ndc80 to plus-ends to prevent kMT detachment and chromosome loss during metaphase and anaphase.

A precise understanding of the mechanisms underlying kMT strengthening via various protein complexes was previously missing due to limitations in spatial and temporal resolutions of current experimental approaches. Here, we developed a new computational model of kMT attachment dynamics incorporating the contributions from Ndc80-associated protein complexes. Our model revealed that the mechanisms of binding, biased diffusion and load-dependent unbinding of Ndc80-associated proteins are sufficient to explain stabilization of kMT attachments. It will be interesting to add a cooperativity factor of protein binding to further understand the roles and synergistic properties of these different protein complexes in stabilizing the kMT interface. In the future, we will incorporate this feature and evaluate how kMT attachments respond to tension, when assuming cooperativity between dynamic proteins. In addition, we will test conditions when Ndc80-associated protein complexes also affect microtubule dynamics. Moreover, it will be interesting in the future to test if the MAPs that assist Ndc80 function form sub-complexes with Ndc80 to provide spatial and temporal control for robust kMT attachment formation. Further, it will be an exciting prospect to probe the nature of structures formed by these complexes on microtubules in an effort to understand how they precisely contribute to these functions. For this purpose, experiments combined with simulations based on all-atom and molecular modeling could help understanding kMT complexes’ structures and functions.

## Materials and Methods

### Cell Culture and transfection

HeLa cells were cultured and maintained in Dulbecco’s modified eagle’s medium (DMEM, Life Technologies) containing 10 % fetal bovine serum (Life Technologies), using standard cell culture procedures (Agarwal et al., 2018). For transfection experiments involving -ch-TOG-GFP, cells were seeded overnight onto 22 mm glass coverslips to achieve ~60-70 % confluency. The cells were then transfected with 0.5 μg of plasmid encoding ch-TOG-GFP (a generous gift from Dr. Stephen Royle at University of Warwick, UK) using Effectene transfection reagent (Qiagen) according to the manufacturer’s instructions. The cells were fixed 48 h post-transfection and processed for immunofluorescence microscopy.

### Cell fixation and immunofluorescence staining

The cells were fixed using ice-cold methanol for 6 mins or 4 % formaldehyde for 20 mins as indicated in the figures or figure legends. For Cdt1 spindle microtubule staining, cells were fixed in ice-cold methanol only for 3 mins. Following fixation, the cells were immune-stained with combinations of different primary antibodies, as indicated in the figures or figure legends. The primary antibodies used in the study include Hec1 monoclonal (1:400, ab3613, clone 9G3, Abcam), Zwint1 polyclonal (1:400, A300-781A, Bethyl), Tubulin monoclonal (1:500, T9026, clone DM1A, Sigma), anti-CREST antiserum (1:500, HCT0100, Immunovision, Inc.) and Cdt1 polyclonal (1:50, H300, Santacruz Biotechnologies). The Rabbit polyclonal antibody against Ska3 used at 1:200 dilution was a kind gift from Dr. Gary Gorbsky (Oklahoma Medical Research Foundation, University of Oklahoma). 4,6-diamino-2-phenylindole (DAPI) dihydrochloride (1:10,000, Life Technologies) was used to counterstain the nucleus/chromosomes. Alexa Fluor 488-, Rhodamine Red-X-, or Cy5-labeled donkey secondary antibodies were obtained from Jackson ImmunoResearch Laboratories, Inc. and used at a dilution of 1:200.

### Spinning-disc confocal microscopy

Following immunostaining, the coverslips were mounted on to glass slides using ProLong Gold Antifade reagent (Invitrogen). For image acquisition, 3D stacks were obtained sequentially at 200nm steps along the z axis through the cell using a high-resolution inverted microscope (Eclipse TiE; Nikon) equipped with a spinning disk (CSU-X1; Yokogawa Corporation of America), an Andor iXon Ultra888 EMCCD camera, and an x100 1.4 NA Plan-Apochromatic DIC oil immersion objective (Nikon). The images were acquired and processed using the NIS elements Software from Nikon.

### Computational model of kMT attachment dynamics

In order to understand the interplay between Ndc80, Ska, and Cdt1 for stabilizing kinetochore-microtubule attachments, we developed a computational model. The use of a computational approach allowed us to directly incorporate the different protein components, isolate their contributions, and characterize their synergistic effects on the emergent dynamics of kMT attachments, including their displacement along tubulin protofilaments and resistance to tension. In the model, Ndc80, Ska1, Cdt1 and ch-TOG are explicitly defined by a position, *x_i_*, on a onedimensional lattice mimicking the tubulin protofilament, and exists in two states, bound or unbound. When in the bound state, each protein moves. It can also unbind depending on its unbinding rate, which determines its unbinding probability. All model parameters, including binding and unbinding rates, diffusion coefficients, and characteristic lifetimes of bound states, are taken from previous experiments.

Initially, all proteins are considered to be at the tip of tubulin protofilaments, at position 0. At each time step of the simulations, each protein can switch between bound and unbound states, and diffuse along the protofilaments; when in the bound state, proteins share tension and undergo biased diffusion; unbound proteins are dragged along by the bound proteins. As a result, during the diffusion along the simulated 1D microtubule lattice, while individual proteins undergo cycles of binding, unbinding and biased diffusion, the kMT interface, formed by all proteins, moves directionally. In summary, the developed algorithm, while updating position and state of each protein individually, micks the directional displacement and force-sensing properties of an integrated kMT attachment interface. As for output, the model is used to evaluate: (*i*) displacement of the kMT interface; (*ii*) time of kMT attachment under tension and (*iii*) kMT attachment rupture force.

### Dynamics of kMT proteins in non-load bearing conditions

We simulated the time evolution of the kMT interface formed by Ndc80, Ska and Cdt1, by developing a computational model based on the Kinetic Monte Carlo approach. According to this approach, a system evolves dynamically from state to state, and the transitions between states are treated directly. At each time step of the simulation, the probability of binding, *P_on_*, is evaluated, based upon the protein binding rate, *k_on_*, as: *P_on_* = *k_on_ dt*. A uniform random number, *u*, is then generated and compared with this probability. If *P_on_* > *u*, then the protein switches to the bound state. Then, for all bound proteins, unbinding is evaluated based upon unbinding probability, *P_off_*, accounting for the unbinding rate, *k_off_*, as: *P_off_* = *k_off_ dt*. Each bound protein undergoes biased diffusion, and its position moves 4 nm, a distance corresponding to the typical size of a tubulin monomer and to Ndc80 spacing within a tubulin protofilament (G. M. Alushin et al., 2010; Zaytsev et al., 2015). In the discrete time and space of our simulation, each bound protein is considered a random walker and it hops to the right with a probability *p* or to the left with a probability (1-*p*), where *p* > *q*, with numerical values chosen to match experimental diffusion rates (Agarwal et al., 2018; Schmidt et al., 2012; Umbreit et al., 2012). However, since the kMT interface moves as a whole, at the end of all time-steps in the simulation, the positions of all proteins are re-assigned. Both unbound and bound proteins are assigned the position of the kMT interface, *x_interface_*, computed based on the average position of all bound proteins, *N_bound_*, as: 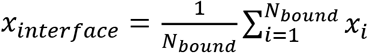. We ran simulations for 300 s, which is a physiologically relevant timescale for kMT attachments (Zaytsev, Ataullakhanov, & Grishchuk, 2013). As for the output, kMT displacement was evaluated.

### Dynamics of kMT proteins in load-bearing conditions

Our model incorporates load-bearing conditions, where tension acts on the kMT attachment. The model assumes that this tension increases at each time step, and modifies the initial protein unbinding rate, *k_off,0_*. Initially, the kMT is unloaded, with null total force, *F_tot_* = 0; then, at each time step, the total force increases randomly and uniformly, with a value in the range 0-0.1 pN. The model assumes that this force is equally distributed on all bound proteins, *n_b_*. Therefore, for each *i* protein in the bound state, the force acting on it is: 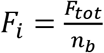. As for the non-load bearing conditions, the basic algorithm evaluates proteins state and position, and updates them at every time step, by evaluating binding, unbinding, and diffusion. Importantly, for the load-bearing conditions, protein unbinding rates are not constant and depend on the load per protein, *F_i_*. For *F_i_* = 0 pN, unbinding rate corresponds to that used in non load-bearing conditions, therefore *k_off,0_* = *k_off_*. For *F_i_* > 0 pN, unbinding rates depend on the type of bond between the protein and microtubules, whether it is a slip bond (as for Ndc80, Ska, and Cdt1) or catch bond (as for ch-TOG). The following session explains how the unbinding rates change with load.

### Slip bond and catch bond dynamics

In slip bonds, bond lifetimes, *τ*, decrease with increasing tension, *F_i_* (Figure 4B). Accordingly, *k_off_* increases under tension. Slip bond dynamics follows a single exponential pathway and the decrease of *τ* with tension is:

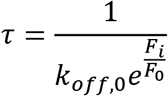

where *k_off,0_* is the unbinding rate at zero force and *F*_0_ = 3 pN for Ndc80 (Civelekoglu-Scholey et al., 2013).

In catch bonds, *τ* initially increases with increasing tension, followed by a decrease (Figure 5B). The resulting bond pathway has both a strengthening and a weakening pathway. Catch bonds are expressed mathematically as double exponentials, where exponents have different signs: a positive sign for the strengthening pathway and a negating one for the weakening pathway. The positive and negative exponential signs correspond to an initial increase in *τ*, followed by a decrease:

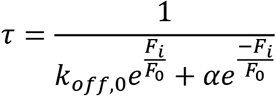

## Acknowledgements

We would like to thank our respective start-up funds from the Department of Bioengineering and the Scientific Computing and Imaging Institute of the University of Utah and the Feinberg School of Medicine, Northwestern University, for supporting our research on this project. This work was also supported by NCI grant R00CA178188 to DV. We would like to thank Drs. Gary Gorbsky and Stephen Royle for help with reagents used in the study.

